# Epicardial Tcf21 facilitates cardiomyocyte dedifferentiation and heart regeneration in zebrafish

**DOI:** 10.1101/2025.05.15.654216

**Authors:** Miglė Kalvaitytė-Repečkė, Sofija Gabrilavičiūtė, Kotryna Kvederavičiūtė, Leonard Burg, Edita Bakūnaitė, Kenneth D. Poss, Darius Balciunas

## Abstract

Unlike mammals, zebrafish (*Danio rerio*) are able to regenerate their hearts after injury, making them an excellent model organism for studying the molecular mechanisms underlying heart regeneration. Epicardium, the outermost layer of the heart, is an essential player in this process. Injury-induced epicardium activation, characterized by the expression of embryonic epicardial marker genes including *tcf21* supports cardiac regeneration by providing various cell types and releasing paracrine signals that promote the restoration of damaged tissue. However, the molecular mechanisms involved in this process are insufficiently understood. In this study, we describe a conditional *tcf21^flox^* allele and use it to investigate the role of Tcf21 in heart regeneration. By employing 4-hydroxytamoxifen inducible CreER^T2^ recombinase, we eliminated *tcf21* expression in adult fish. Our findings indicate that loss of this transcription factor reduces the presence of dedifferentiated cardiomyocytes in the injury area and impairs heart regeneration. This work provides new insights into the molecular basis of the epicardial response to heart injury and its role in guiding heart regeneration.

## INTRODUCTION

Cardiovascular diseases remain the leading cause of death worldwide, with coronary heart disease that often leads to myocardial infarction being the most prevalent form (Jayaraj et al., 2018; Martin et al., 2024). Adult mammals have limited capacity to replace damaged cardiomyocytes, leaving the injured heart unable to recover full functionality (Bergmann et al., 2009; Broughton et al., 2018). In contrast, zebrafish (*Danio rerio*) possess the remarkable ability to regenerate their hearts throughout their lives (Poss et al., 2002). Following the initial response to the injury (González-Rosa et al., 2011; Lodrini & Goumans, 2021; Sun et al., 2002), zebrafish cardiomyocytes undergo dedifferentiation and proliferate, generating new myocardium to repair the damaged tissue (Jopling et al., 2010; Kikuchi et al., 2010).

Studies have highlighted the importance of the epicardium, the outermost layer of the heart, in supporting heart regeneration (reviewed in (Cao & Poss, 2018; Quijada et al., 2020; Simões & Riley, 2018)). Initial pan-epicardial activation, characterized by the upregulation of embryonic epicardium marker genes like *tbx18, wt1b*, and *aldh1a2*, and proliferation, becomes localized to the injury area by 7 days post-injury (Kikuchi, Gupta, et al., 2011; Kikuchi, Holdway, et al., 2011; Lepilina et al., 2006; Schnabel et al., 2011; van Wijk et al., 2012). During regeneration, the epicardium secretes various signaling molecules and provides diverse cell types supporting extracellular matrix (ECM) remodeling, cardiomyocyte proliferation, and neovascularization of the damaged area (Allanki et al., 2021; Fang et al., 2013; González-Rosa et al., 2011; Lepilina et al., 2006; Sánchez-Iranzo et al., 2018; J. Wang et al., 2013, 2015).

Tcf21 (transcription factor 21, also known as Pod-1, Capsulin, or Epicardin) plays an essential role in heart development and epicardium formation (Tandon et al., 2013). Its loss leads to neonatal lethality in mice (Braitsch et al., 2012; Lu et al., 2000; Quaggin et al., 1999), while zebrafish larvae display severe defects in craniofacial muscle formation and cardiac chamber morphogenesis that eventually lead to death (Boezio et al., 2023; Burg et al., 2016; Nagelberg et al., 2015). This transcription factor is expressed in the majority of the epicardial cells and regulates the differentiation of epicardium-derived cells (EPDCs) into cardiac fibroblasts and smooth muscle cells (Acharya et al., 2012; Braitsch et al., 2012; Lu et al., 1998; Weinberger et al., 2020). Tcf21 expression persists in the dormant epicardium and resting fibroblasts in the adult heart (Braitsch et al., 2013; Kanisicak et al., 2016; Kikuchi, Gupta, et al., 2011; Lepilina et al., 2006; Weinberger et al., 2024). However, the role of Tcf21 in heart regeneration remains unexplored, primarily because of the challenge of conditionally inactivating *tcf21* in adult zebrafish.

Recent advances in genome editing enabled the engineering of conditional mutants, allowing to study gene function in adult fish (Kalvaitytė & Balciunas, 2022). In this study, we generated a conditional (floxed) *tcf21* allele to investigate the role of Tcf21 in heart regeneration. Using globally expressed 4-hydroxytamoxifen (4-HT) inducible CreER^T2^ recombinase, we disrupted *tcf21* expression in adult fish and found that the loss of Tcf21 impairs heart regeneration. The analysis of organ-wide transcriptome following the injury indicated higher expression of sarcomere-related genes and downregulation of several chemokines. Further investigation uncovered compromised dedifferentiation of cardiomyocytes in the injury area, highlighting the role of epicardial Tcf21 in creating a permissive environment for heart regeneration.

## RESULTS AND DISCUSSION

### *tcf21^flox^* allele can be conditionally disrupted during zebrafish development

To study the role of *tcf21* in heart regeneration, we generated a conditional *tcf21* floxed allele by flanking the first exon with two directly oriented *loxP* sites. We previously reported the *tcf21^tpl144^* allele, which has a *loxP* site integrated into the 5’ UTR (Burg et al., 2018). Following the same methodology, we engineered the second *loxP* site into the intron (**Fig. 1A; Fig. S1A,B**). A single F1 fish with precise integration of both *loxP* sites was selected to establish a stable transgenic *tcf21^vln18^* (*tcf21^flox^*) zebrafish line (**Fig. S1C,D**). Genotyping offspring of the heterozygous *tcf21^flox^* fish incross confirmed that homozygous *tcf21^flox/flox^* fish were viable and indistinguishable from their wild-type counterparts (**Fig. S1E,F**).

**Fig. 1.**
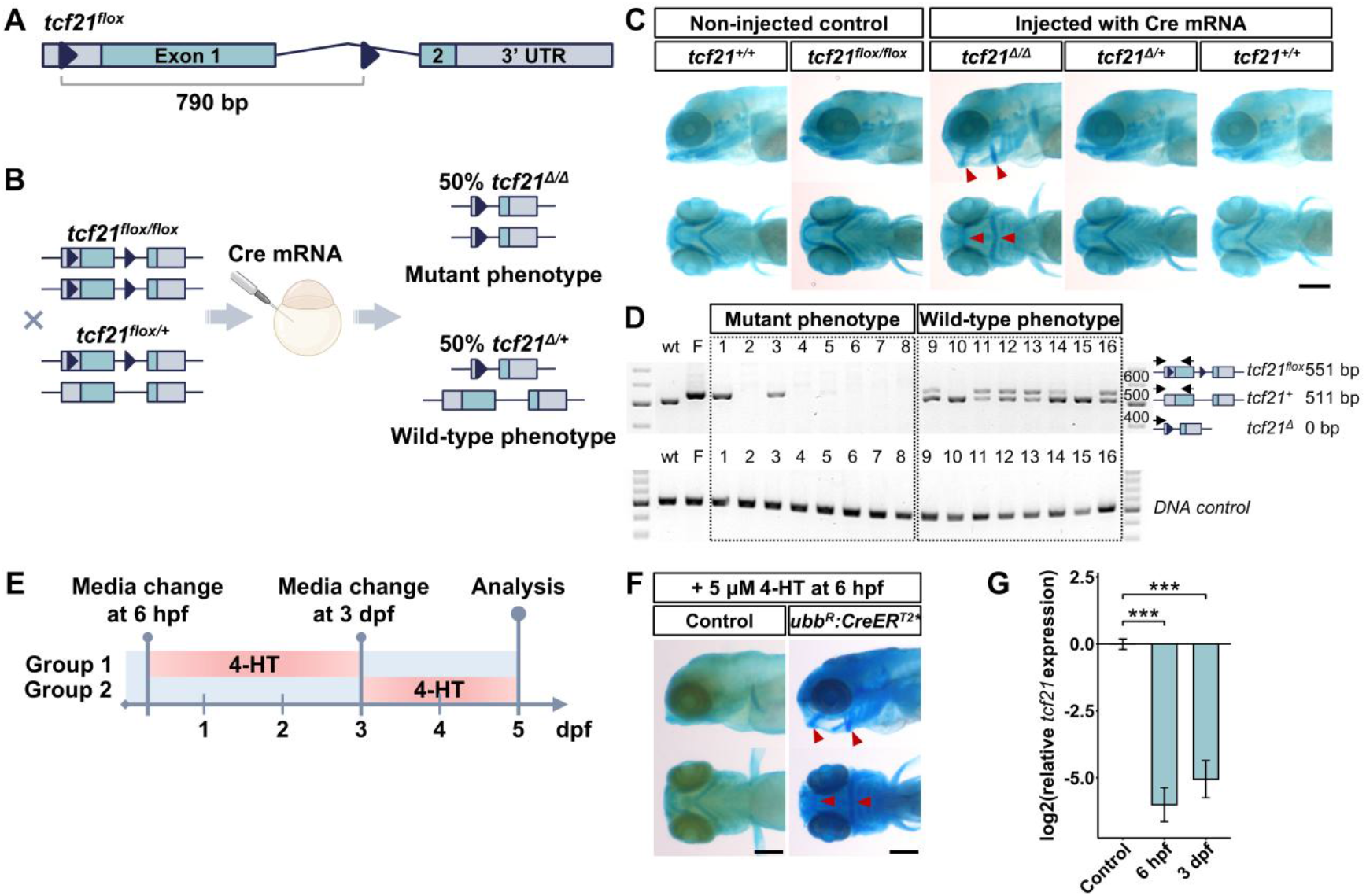
Validation of the *tcf21^flox^* allele. **A.** Diagram of the engineered *tcf21^flox^* allele. Triangles mark *loxP* sites. **B.** Experimental design. **C.** 5 dpf larvae stained with Alcian blue. **D.** Genotyping of individual larvae displaying mutant or wild-type phenotypes from (**C**). “wt” – *tcf21^+/+^* control; “F” – *tcf21^flox/flox^* control. **E.** Experimental timeline of *tcf21^flox/flox^, ubb^R^:CreER^T2^\** embryo treatment with 5 μM 4-HT. **F.** 5 dpf *tcf21^flox/flox^* (control) and *tcf21^flox/flox^, ubb^R^:CreER^T2^\** larvae treated with 4-HT from 6 hpf to 3 dpf. **G.** *tcf21* expression in 5 dpf *tcf21^flox/flox^, ubb^R^:CreER^T2^\** larvae treated with 4-HT from 6 hpf to 3 dpf or from 3 dpf to 5 dpf. Results were normalized to the *tcf21* expression in *tcf21^flox/flox^* larvae, treated with 4-HT from 6 hpf to 3 dpf, and transformed to a logarithmic scale. Data are represented as mean±SD. *P* value was calculated using one-way ANOVA followed by Tukey’s post-hoc test; \*\*\**P*<0.001. Red arrowheads in (**C,F**) mark defects in branchial arch formation; scale bar: 200 μm.

To validate the engineered allele, we crossed homozygous and heterozygous *tcf21^flox^* fish and injected single-cell-stage embryos with Cre recombinase mRNA (Balciuniene et al., 2013; Burg et al., 2018) (**Fig. 1B**). After successful recombination, a 790 bp DNA fragment encoding the DNA-binding domain is expected to be excised, resulting in *tcf21* knock-out (*tcf21^Δ^*). Around 40% of Cre-injected embryos displayed severe facial malformations at 5 dpf, consistent with the previously documented loss-of-function phenotype of homozygous *tcf21* mutants (**Fig. 1C**) (Burg et al., 2016; Lee et al., 2011; Nagelberg et al., 2015). Genotyping confirmed that larvae with severe branchial arch defects were initially homozygous for the *tcf21^flox^* allele (**Fig. 1D**). Meanwhile, heterozygous siblings retained wild-type phenotype, confirming that the generated *tcf21^flox^* allele can be conditionally disrupted using Cre recombinase.

The usage of conditional alleles relies on the efficiency of the Cre-recombinase-expressing driver line. We assessed the conditional disruption of the *tcf21^flox^* gene using 4-HT-inducible epicardium-specific *TgBAC(tcf21:CreER^T2^)pd42 (tcf21:CreER)* transgene (Kikuchi, Gupta, et al., 2011), which was used to conditionally knock out *shha*^*ct*^ allele (*Tg(shha:Zwitch)vcc8Gt*) in zebrafish larvae and adults (Sugimoto et al., 2017). We treated *tcf21^flox/flox^, tcf21:CreER* embryos from 6 to 72 hpf with different concentrations of 4-HT ranging from 5 to 15 μM and analyzed the phenotype of the larvae at 5 dpf by Alcian blue staining. Unexpectedly, none of the larvae displayed phenotypes consistent with complete loss of function (**Fig. S2A**). To rule out the possibility of poor activity of 4-HT (Felker et al., 2016), we used *Tg(−3.5ubb:loxP-EGFP-loxP-mCherry)cz1701 (ubi:Switch)* reporter line (Mosimann et al., 2011), and confirmed that 5 μM 4-HT treatment from 6 to 72 hpf activates the recombinase, changing the fluorescent signal from green to red in *tcf21+* cells (**Fig. S2B,C**). Therefore, we concluded that the *tcf21:CreER* transgene does not achieve complete conditional knock-out of the *tcf21^flox^* allele and is unsuitable for further analysis.

A recently described *Tg(ubb^R^:CreER^T2^*)vln2 (ubb^R^:CreER^T2^\**) transgenic line (Bakūnaitė et al., 2024) displays high recombination efficiency throughout development and adulthood; thus, we tested its ability to knock out the *tcf21^flox^* allele. We treated *tcf21^flox/flox^, ubb^R^:CreER^T2^\** embryos from 6 to 72 hpf with 5 μM 4-HT and observed loss-of-function phenotype and nearly complete loss of *tcf21* expression in 5 dpf larvae (log_2_(*tcf21*) = -6.01±0.63 (1.7±0.8%), *P*=0.0008) (**Fig. 1E-G; Fig. S2D,E**). Similar results were obtained by treating larvae from 3 to 5 dpf (log_2_(*tcf21*) = -5.05±0.70 (3.3±0.2%), *P*=0.00043), confirming the effective disruption of the *tcf21^flox^* allele during development (**Fig. 1E,G; Fig. S2F**).

### Conditional *tcf21* knock-out in adult fish leads to impaired heart regeneration

Although several conditional zebrafish mutants have been created using various methodologies, only a handful have been applied in adult fish (Angom et al., 2023; Grajevskaja et al., 2018; Ogawa et al., 2021; Rajan et al., 2024; Sugimoto et al., 2017). After successful inactivation of the *tcf21^flox^* allele at later developmental stages using *ubb*^*R*^:*CreER*^*T2*^* transgene, we assessed recombination efficiency in adult fish. 3 to 12-month-old *tcf21^flox/flox^, ubb^R^:CreER^T2^\** fish were immersed into 5 μM 4-HT solution three times for 24 h, allowing them to recover for 24 h at normal husbandry conditions between each treatment. After the third treatment, fish were allowed to recover for a week (**Fig. 2A**). We collected ventricles for RNA and DNA extraction at 0 days post-treatment (dpt) without injury and 3 days post-cryoinjury (dpci) to assess changes in *tcf21* expression during regeneration. We used untreated wild-type and *tcf21^flox/flox^, ubb^R^:CreER^T2^\** fish as controls and collected their ventricles without injury or at 3 dpci. RT-qPCR analysis showed nearly three-fold upregulation of *tcf21* expression at 3 dpci in untreated fish, while the amount of *tcf21* mRNA was significantly reduced to almost undetectable level in 4-HT-treated *tcf21^flox/flox^, ubb^R^:CreER^T2^\** fish ventricles both at 0 dpt (log_2_(*tcf21*) = -6.04±0.51 (1.6±0.5%), *P*=0.002) and 3 dpci (log_2_(*tcf21*) = -5.39±0.97 (2.8±1.5%), *P*=0.002)(**Fig. 2B)**. Moreover, no difference in *tcf21* expression was detected between uninjured wild-type and uninjured, untreated *tcf21^flox/flox^, ubb^R^:CreER^T2^\** fish, confirming that integrated *loxP* sites do not perturb *tcf21* expression in the adult fish heart.

**Fig. 2.**
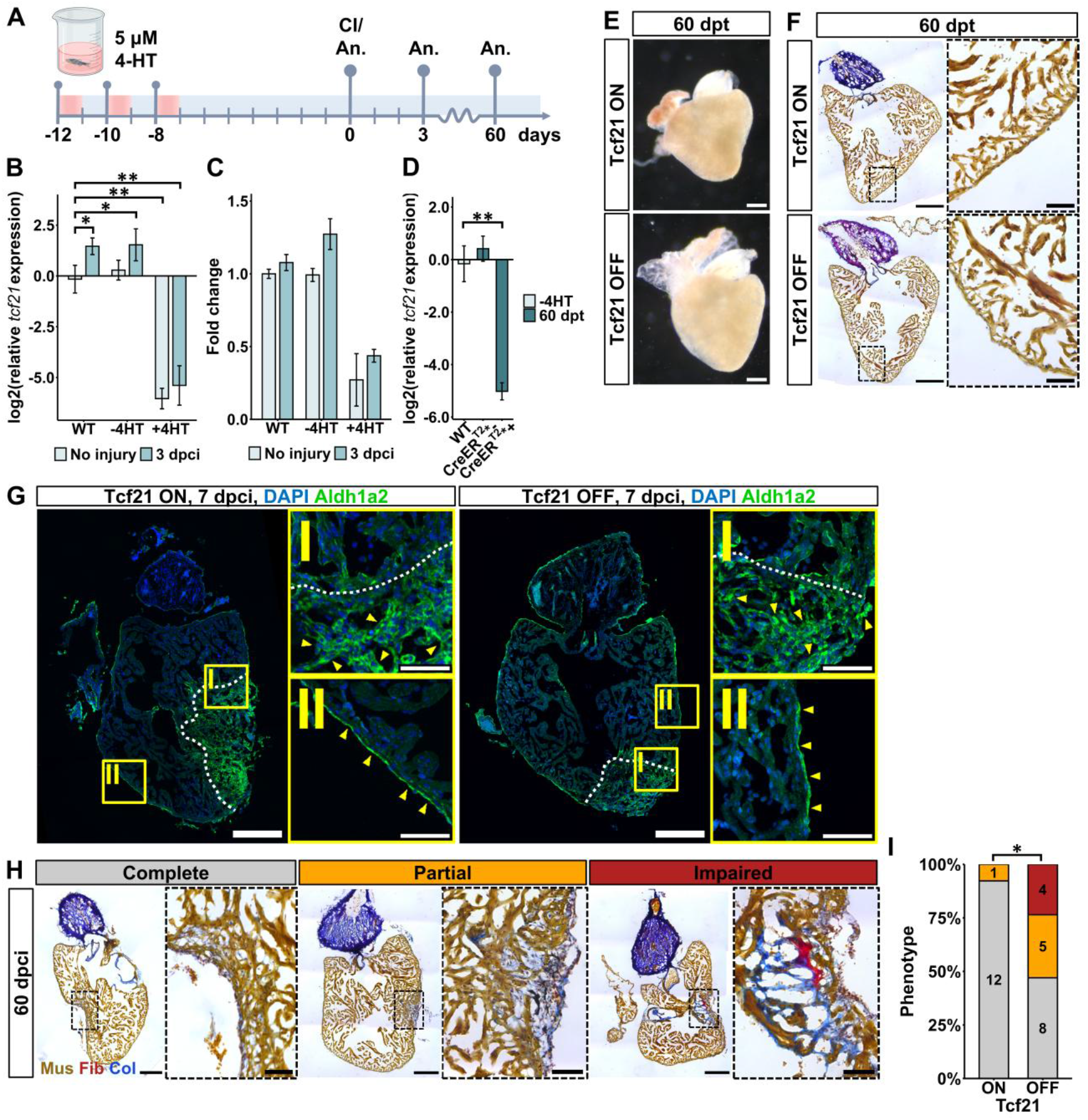
*tcf21^flox^* knock-out in adult fish impairs heart regeneration. **A.** Experimental timeline. “CI” – cryoinjury; “An.” – analysis. **B-C.** *tcf21* expression (**B**) and *tcf21* knock-out at the gDNA level (**C**) in ventricles of wild-type (WT), untreated (−4HT), and 4-HT-treated (+4HT) *tcf21^flox/flox^, ubb^R^:CreER^T2^\** fish at 0 dpt (n=3) and 3 dpci (n=3). **D.** *tcf21* expression in ventricles of 4-HT-treated *tcf21^flox/flox^* (CreER^T2^*-) and *tcf21^flox/flox^, ubb^R^:CreER^T2^\** (CreER^T2^*+) fish at 60 dpt (n=3). Results in (**B-D**) were normalized to the wild-type fish samples at 0 dpt; *tcf21* expression data in (**B,D**) was transformed to a logarithmic scale. Data are shown as mean±SD. **E-F.** Bright-field images (**E**) and AFOG-stained heart sections (**F**) of adult fish hearts. Black boxes in (**F**) mark the areas shown in zoomed images. **G.** Aldh1a2 staining of heart sections. White dashed lines indicate wound border, yellow boxes mark the areas shown in zoomed images, and yellow arrowheads point to Aldh1a2-expressing epicardial cells. **H.** Representative sections of the AFOG-stained hearts from each category with zoomed images of the marked area. **I.** Diagram showing the distribution of regenerative phenotypes from (**H**). *P* value in (**B-D**) was calculated using the pairwise Wilcoxon rank-sum test with Bonferroni correction; in (**I**) using Fisher’s Exact Test; \**P*<0.05, \*\**P*<0.01, \*\*\**P*<0.001. Scale bars: 200 μm in (**E-H**); 50 μm in (**F-H**) zoomed images.

We also performed qPCR to determine knock-out efficiency at the DNA level in *tcf21^flox/flox^, ubb^R^:CreER^T2^\** fish. After 4-HT-induced recombination, approximately 30-40% of the full-length *tcf21* gene remained at 0 dpt and 3 dpci (**Fig. 2C)**. As the analysis was performed on whole ventricle DNA, we hypothesize that recombination was inefficient in *tcf21*-non-expressing cells where the *tcf21^flox^* gene is not accessible to Cre recombinase due to being packed in heterochromatin. To test if the expression of *tcf21* may recover within the experimental time frame, we analyzed the level of *tcf21* mRNA in 4-HT-treated *tcf21^flox/flox^, ubb^R^:CreER^T2^\** fish at 60 dpt. We confirmed a significant reduction of *tcf21* mRNA (log_2_(*tcf21*) = -5.02±0.33 (3.1±0.7%), *P*=0.009), demonstrating that there is almost no detectable full-length *tcf21* mRNA capable of producing functional Tcf21. Thus, henceforth, we refer to 4-HT-treated *tcf21^flox/flox^, ubb^R^:CreER^T2^\** fish as “Tcf21 OFF”, while 4-HT-treated *tcf21^flox/flox^* or untreated *tcf21^flox/flox^, ubb^R^:CreER^T2^\** fish are “Tcf21 ON”.

Previous studies showed that *Tcf21* null embryos in mice exhibit epicardium blistering from stage E14.5 (Acharya et al., 2012; Braitsch et al., 2012). Meanwhile, fibroblast-specific inactivation of *Tcf21^fl/fl^* in adult mouse hearts did not alter the homeostasis of cardiac fibroblasts, nor their function in promoting cardiac fibrosis (Johansen et al., 2025). We compared the phenotypes of Tcf21 ON and Tcf21 OFF fish hearts at 60 dpt and found no significant differences (**Fig. 2E,F**). We next sought to investigate if loss of Tcf21 would affect heart regeneration. We treated *tcf21^flox/flox^* and *tcf21^flox/flox^, ubb^R^:CreER^T2^\** fish with 4-HT and analyzed heart sections at 7 and 60 dpci. Epicardial cells were present in the injury area of both Tcf21 ON and Tcf21 OFF hearts at 7 dpci, as indicated by immunostaining for Aldh1a2 (**Fig. 2G**) (Kikuchi, Holdway, et al., 2011; J. Wang et al., 2011). Myocardial regeneration of hearts at 60 dpci was scored blindly from AFOG-stained histological sections by an independent expert as completely regenerated, partially regenerated, and impaired (**Fig. 2H,I; Fig. S3**). Strikingly, over half of injured Tcf21 OFF ventricles (9 out of 17) displayed partially or fully blocked regeneration of the myocardial wall with an increased amount of collagen and fibrin present in the injury area (*P*=0.0395). In contrast, consistent with previous observations, almost all control Tcf21 ON fish (12 out of 13) regenerated a contiguous myocardium wall during that time (Chablais et al., 2011; González-Rosa et al., 2011). The increased amount of connective tissue and the lack of newly formed myocardium that covers the injury area allows us to conclude that the loss of Tcf21 impairs heart regeneration.

### Whole-ventricle RNA-seq analysis suggests a role for Tcf21 in promoting cardiomyocyte dedifferentiation

Following heart injury, dormant epicardium cells are activated to divide and cover the area of trauma. The organ-wide response peaks at 3 dpi and becomes restricted to the injury site by 7 dpi (González-Rosa et al., 2011; Kikuchi, Gupta, et al., 2011; Kikuchi, Holdway, et al., 2011; Lepilina et al., 2006; Schnabel et al., 2011; J. Wang et al., 2013). To analyze the early regenerative changes in gene expression, we performed bulk RNA-seq on whole ventricle mRNA of Tcf21 ON and Tcf21 OFF fish without injury (D0), 3 dpci (C3), and 3 days post-sham (dps) injury (S3) (**Fig. 3A**). Principal-component analysis (PCA) showed little gene expression variability between uninjured hearts, which became more distinct at 3 dpci and 3 dps (PC1, 47%) (**Fig. 3B)**. The analysis of differentially expressed genes (DEGs) (*P*_adj_<0.05) between different conditions indicated 1,011 unique genes that were differentially expressed in Tcf21 OFF hearts following the cryoinjury (**Fig. 3C**). Out of those, several genes with roles in cell adhesion, including *col28a1b, ctnna2, thbs1b,* and *podxl*, also associated with the epithelial identity of the epicardium, were upregulated (Gebauer et al., 2016; Shen et al., 2018; Vite et al., 2015; Weinberger et al., 2020; Xia et al., 2022).

**Fig. 3.**
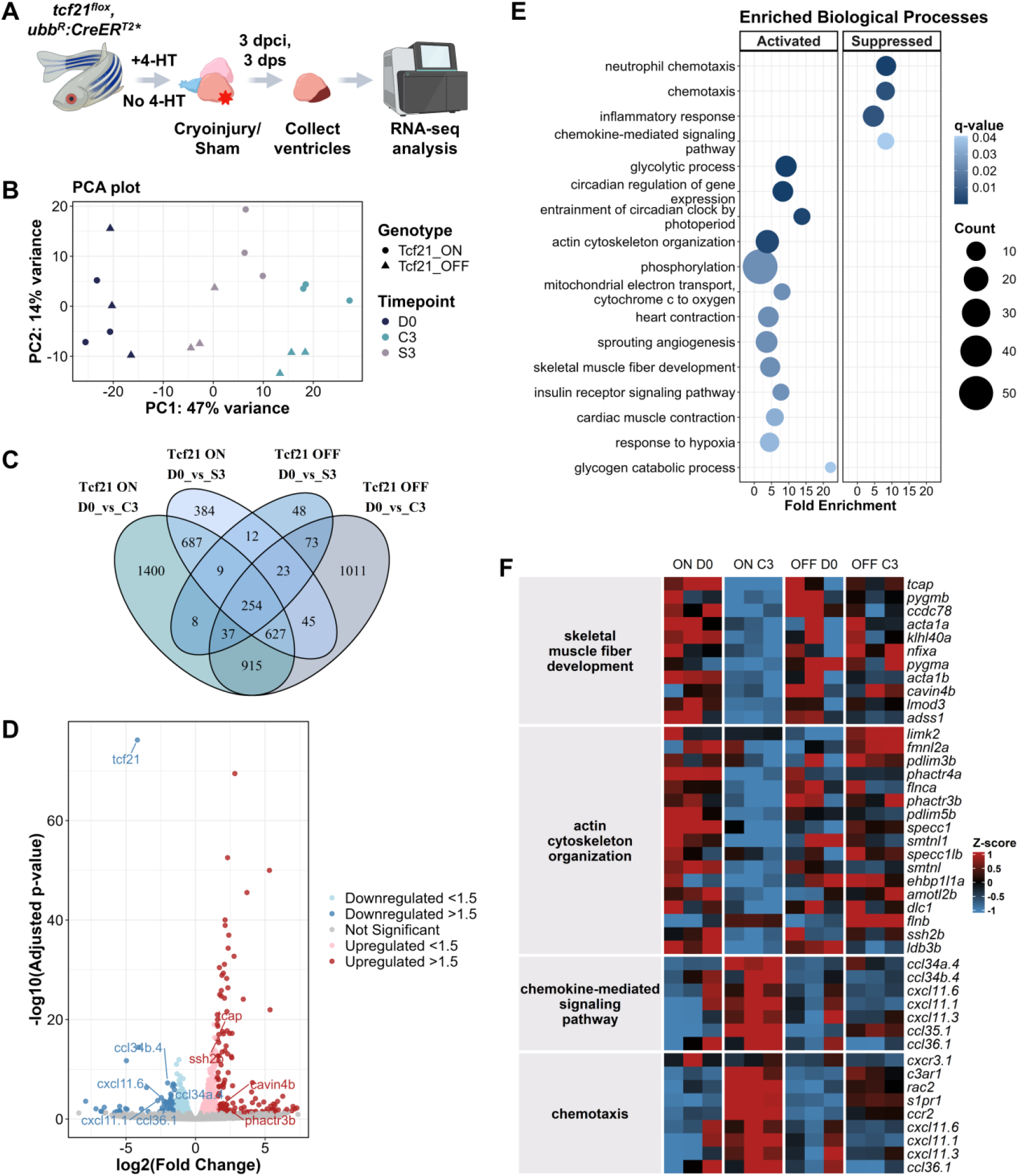
Whole ventricle RNA-seq indicates a potential Tcf21 role in promoting cardiomyocyte dedifferentiation during heart regeneration. **A.** Experimental design. 4-HT-treated (Tcf21 OFF) and untreated (Tcf21 ON) *tcf21^flox/flox^, ubb^R^:CreER^T2^\** fish were used to collect ventricles for bulk RNA-seq at 0 dpt (D0), 3 dpci (C3), and 3 dps (S3). **B.** PCA of transcriptome samples. **C.** Venn diagram depicting the overlap of DEGs across conditions. **D.** Volcano plot representing DEGs at 3 dpci in Tcf21 ON and Tcf21 OFF hearts. **E.** Enriched biological processes GO terms of significantly downregulated and upregulated genes at 3 dpci. **F.** Expression of genes from selected GO terms during regeneration in Tcf21 ON and Tcf21 OFF hearts.

A total of 1,175 DEGs (*P*_adj_<0.05) were found at 3 dpci by comparing gene expression in Tcf21 ON versus Tcf21 OFF hearts, of which 375 genes were downregulated and 800 – upregulated (**Fig. 3D**). Among downregulated genes, there were several chemokine ligands, such as *ccl34a.4, ccl34b.4, cxcl11.6, cxcl11.1, ccl36.1;* meanwhile, upregulated genes were related to actin cytoskeleton, such as *tcap* and *cavin4b,* which localize to Z-disc and are involved in T-tubule organization (Ben-Yair et al., 2019; Housley et al., 2016; Münch et al., 2017; Zhang et al., 2009; Zhou et al., 2020). Gene enrichment analysis indicated that downregulated genes were associated with “chemotaxis” (q-value=0.006), “chemokine-mediated signaling pathway” (q-value=0.04), and other related biological processes. Meanwhile, upregulated genes were associated with sarcomere-related GO terms, such as “actin cytoskeleton organization” (q-value=0.003), “skeletal muscle fiber development” (q-value=0.02), and “heart contraction” (q-value=0.02) (**Fig. 3E**). We selected several GO terms and analyzed the expression dynamics of assigned genes during the regeneration. Our analysis revealed that in Tcf21 OFF hearts, expression of these genes mainly remained unaffected, while in control hearts, the changes were significant (**Fig. 3F**). These results indicate that Tcf21 function in epicardial cells contributes to suppressing actin cytoskeleton organization and cardiac muscle fiber development while activating the expression of various chemokines involved in chemotaxis during heart regeneration.

### Loss of Tcf21 inhibits cardiomyocyte protrusion into the injured area

Heart regeneration occurs through cardiomyocyte (CM) dedifferentiation and proliferation, with some studies suggesting displacement or active migration into the injury area from the border zone (Beisaw et al., 2020; Ben-Yair et al., 2019; Constanty et al., 2024; Gemberling et al., 2015; Itou et al., 2012; Jopling et al., 2010; Kikuchi et al., 2010; Marín-Juez et al., 2019; Morikawa et al., 2015; J. Wang et al., 2011; Wu et al., 2016). RNA-seq results indicated that in Tcf21 OFF hearts, sarcomere disassembly and CM dedifferentiation are compromised, potentially hindering muscle restoration. We assessed CM dedifferentiation by immunostaining for embCMHC at 7 dpci (Ben-Yair et al., 2019; Pfefferli & Jaźwińska, 2017; Sallin et al., 2015). We observed fewer dedifferentiated CM in the injury area in Tcf21 OFF hearts compared to the control (*P*=0.0056) (**Fig. 4A,B**). Interestingly, while the embCMHC signal was present in both conditions, the number (*P*=0.0022) and length (*P*=5.055986e-40) of CM protrusions into the injured area were significantly reduced in Tcf21 OFF hearts (**Fig. 4C,D**). Notably, proliferation of border zone CMs showed no significant difference between Tcf21 ON and Tcf21 OFF hearts at 3 or 7 dpci (**Fig. 4E-G; Fig. S4**). This data indicates that epicardial Tcf21 facilitates CM dedifferentiation and normal muscle regeneration following cryoinjury.

**Fig. 4.**
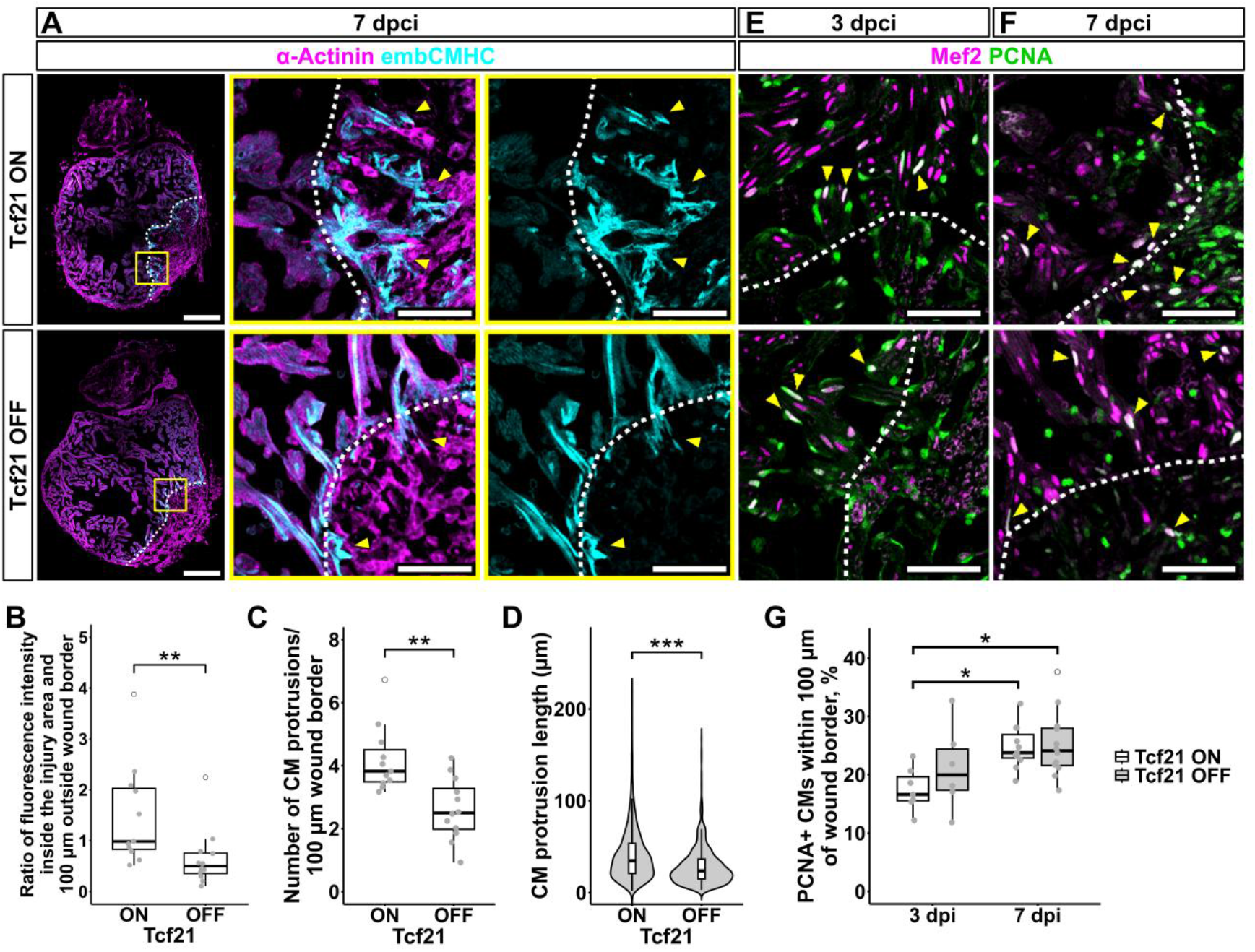
Cardiomyocyte (CM) protrusions into the injured area are disrupted in conditional *tcf21* knock-outs. **A.** Representative heart sections stained for α-Actinin and embCMHC. Yellow arrowheads point to CM protrusions; yellow boxes mark the areas shown in zoomed images. **B.** Quantification of the embCMHC fluorescence intensity ratio in the injury area and 100 μm outside the wound border in Tcf21 ON (n=11) and Tcf21 OFF (n=12) hearts at 7 dpci. **C.** The number of CM protrusions per 100 μm of wound border at 7 dpci. **D.** Distribution of CM protrusion length from the wound border into the injury at 7 dpci. **E-F.** PCNA and Mef2 staining of heart sections at 3 dpci (**E**) and 7 dpci (**F**). Yellow arrowheads point to proliferating (PCNA+) CMs. **G.** Quantification of the PCNA+ CMs compared to the total number of CMs within 100 μm of the wound border in Tcf21 ON (n=6) and Tcf21 OFF (n=6) hearts at 3 dpci, and Tcf21 ON (n=11) and Tcf21 OFF (n=12) hearts at 7 dpci. *P* value in (**B, C**) was calculated using the Wilcoxon rank-sum test; in (**D**) using the unpaired t-test; in (**G**) using one-way ANOVA followed by Tukey’s post-hoc test; \**P*<0.05, \*\**P*<0.01, \*\*\**P*<0.001. Each data point in (**B, C, G**) represents the average of at least two measured heart sections per ventricle. Hollow dots mark outliers. White dashed lines in (**A, E, F**) indicate wound border. Scale bars: 200 μm in (**A**); 50 μm in (**E,F**) and (**A**) zoomed images.

In conclusion, the *tcf21^flox^* allele provides opportunities to investigate gene function in adult fish. Our findings underscore the important role of Tcf21 in heart regeneration, highlighting the significance of epicardium-myocardium crosstalk. The reduced number of dedifferentiated cardiomyocytes in the injury area may reflect changes in EPDCs, ECM composition, or paracrine signaling, all vital for a regenerative environment (J. Wang et al., 2013). Further experiments, such as single-cell transcriptomics or lineage tracing, could clarify how Tcf21 influences the interaction between different cardiac cells during regeneration. This work provides new insights into the molecular basis of cardiac repair and opens new avenues for understanding how transcriptional regulators orchestrate regenerative processes in complex tissues.

## MATERIALS AND METHODS

### Zebrafish maintenance

Wild-type and transgenic zebrafish (*Danio rerio*) were used in accordance with Temple University Institutional Animal Care and Use Committee (IACUC) guidelines under the approval from protocol numbers ACUP 4354, ACUP 4709, and/or in accordance with an approved license of Animal research ethics committee (Lithuania) No. G2-231. Male and female breeders aged 3-18 months were used to generate fish for all experiments. Transgenic zebrafish lines used in this study: *tcf21^tpl144^* (Burg et al., 2018), *tcf21^vln18^* (this study), *TgBAC(tcf21:DsRed2)pd37* (Kikuchi, Gupta, et al., 2011), *TgBAC(tcf21:CreER^T2^)pd42* (Kikuchi, Gupta, et al., 2011), *Tg(−3.5ubb:loxP-EGFP-loxP-mCherry)cz1701* (Mosimann et al., 2011), *Tg(ubb^R^:CreER^T2^*)vln2* (Bakūnaitė et al., 2024). Embryos and larvae were anesthetized with 4 mg/ml MS-222 (Sigma-Aldrich, E10521) dissolved in ddH_2_O and diluted in egg water for handling when necessary.

### Generation of conditional *tcf21^flox^* allele

Integration of the second *loxP* site into the first intron of the *tcf21^tpl144^* allele and identification of the *tcf21^flox^* (*tcf21^vln18^*) allele was performed as previously described (Burg et al., 2018). Briefly, nCas9n mRNA was synthesized from linearized pT3TS-nCas9n (Jao et al., 2013) using the T3 mMESSAGE mMACHINE™ *in vitro* transcription kit (Invitrogen, AM1348). Transcribed mRNA was purified using the RNeasy MinElute kit (Qiagen, 74204), diluted to 150 ng/μL in Nuclease-Free Water, and 2 μL aliquots were stored at -80 °C. sgRNA was prepared using the cloning-free PCR method, diluted to approximately 60 ng/μL, and 8 μL aliquots were stored at -80 °C. Aliquots of sgRNA and nCas9n mRNA were mixed, and 3 nL of the mix were injected into the yolks of single-cell stage embryos obtained from a cross between *tpl144* heterozygote and homozygote, followed by injection of 1 nL of 50 ng/μL *loxP* HDR oligonucleotide as described previously (Burg et al., 2016, 2018). Initial testing of *loxP* integration was performed by PCR on pools of 10-20 injected embryos with multiple combinations of flanking and *loxP*-specific primers. Siblings of successfully injected embryos were raised to adulthood and screened for germline transmission by either out- or in-crossing fish and screening 3 pools of 20 embryos by nested PCR as previously described (Burg et al., 2018). Adult F1 fish were genotyped by tail clip. *tcf21* locus was amplified, and the integration of the full-length *loxP* site was confirmed by Sanger sequencing. Single F1 fish with precise integration of the second *loxP* site into loxP-containing chromosome was used as a founder of the *tcf21^flox^* (*tcf21^vln18^*) fish line. Sequences of HDR oligonucleotide and primers used in this study are listed in Table S1.

### Cre-mediated excision of *tcf21^flox^* allele

Cre mRNA was *in vitro* transcribed using an XbaI-linearized pT3TS-Cre (pDB638) template (Balciuniene et al., 2013) and T3 mMESSAGE mMACHINE™ *in vitro* transcription kit (Invitrogen, AM1348). Transcribed mRNA was purified using RNeasy MinElute Kit (Qiagen, 74204) and diluted to 40 ng/μL in RNase-free water. 2 μL aliquots were stored at -80 °C. Homozygous and heterozygous *tcf21^flox^* fish were crossed, and single-cell-stage embryos were injected with 25 pg of Cre mRNA as previously described (Balciuniene & Balciunas, 2013; Burg et al., 2018). Embryos were analyzed by Alcian blue staining at 5 dpf.

### Alcian blue staining

Embryos were treated with 0.003% PTU (Acros Organics, 207250250) dissolved in DMSO at 12-24 hpf, and the medium was replaced daily until the larvae reached 5 dpf. At 5 dpf, larvae were euthanized by MS-222 (Sigma-Aldrich, E10521) overdose and transferred to a 1.5 mL tube. The specimens were washed twice with 0.1% Tween 20 in PBS (PBST). Then, larvae were fixed with 4% PFA for 4 h at room temperature (RT) on the rocking shaker. Fixative was removed, and the larvae were washed 4 times for 30 min with 0.1% PBST. Larvae were incubated in Alcian blue solution (7.5 mg of Alcian Blue dissolved in 10 mL of acetic acid and 40 mL of ethanol) overnight at RT on the rocking shaker and rehydrated through a graded series of alcohols (100%, 80%, 60%, 40%, and 20% of ethanol in 0.1% PBST) to 0.1% PBST by 1-hour incubation in each solution on the rocking shaker at RT. Finally, larvae were washed twice with 0.1% PBST and mounted in methylcellulose for imaging using a stereo microscope ZEISS Stemi 305 (ZEISS). After imaging, individual larvae were placed in a PCR tube, washed with PBS, and lysed for genotyping. Primers flanking 5’ UTR *loxP* site (tcf21_5inter_F2 and tcf21-R5) were used to identify *tcf21^flox^/ tcf21^+^* alleles. Tbx5a-specific primers were used for DNA control. Sequences of primers are listed in Table S1.

### 4-HT treatment

5 mM 4-HT stock was prepared by dissolving 25 mg of (Z)-4-Hydroxytamoxifen (Sigma-Aldrich, H7904) in 12.9 mL of 96% ethanol by vortexing for 15 min, aliquoted and stored at -80 °C. Before handling, the aliquot was heated for 10 min at 65 °C (Felker et al., 2016). For embryo treatment, up to 60 embryos per petri dish were placed in egg water with 4-HT added to a final concentration of 5 μM at 6 hpf and kept until 3 dpf unless stated otherwise. For treatment of adult fish, individual fish were placed in 100 mL of 5 μM 4-HT solution three times for 24 h in the dark, allowing them to recover for 24 h at normal husbandry conditions between each treatment. After the third treatment, fish were allowed to recover for a week before further procedures.

### Cryoinjury

Zebrafish of 4-12 months of age were used for cryoinjury as described previously (Chablais et al., 2011; González-Rosa et al., 2011). Briefly, adult fish were anesthetized by immersion into 0.2 mg/mL MS-222 (Sigma-Aldrich, E10521) and immobilized with the ventral side upwards in a foam holder mounted on a petri dish. The pericardial sac was opened by a small incision to expose the heart. The apex of the ventricle was touched with a liquid-nitrogen-cooled cryoprobe until the probe was fully thawed. After that, the fish were transferred to a larger container with system water for recovery. Exposing the ventricle without injury was performed for sham controls. Cryoinjured and sham control hearts were harvested at indicated time points after cryoinjury/sham.

### RT-qPCR

RNA was extracted from 25 5 dpf larvae or one adult ventricle per sample using TRI Reagent (Sigma-Aldrich, T9424) and treated with DNase I (Thermo Scientific, 89836) according to the manufacturers’ recommendations. The concentration of RNA was evaluated using a NanoDrop 2000 spectrophotometer (Thermo Scientific). 2 µg of RNA extracted from 5 dpf larvae or 300-400 ng of RNA extracted from adult ventricle was used for cDNA synthesis using Maxima H Minus cDNA Synthesis Master Mix (Thermo Scientific, M1662) following manufacturer’s recommendations. RT-qPCR was performed using 10 ng of cDNA, 0.15 µM of forward and reverse primers, and Maxima SYBR Green qPCR Master Mix (2x) without ROX passive dye (Thermo Scientific, K0253). RT-qPCR was performed on Rotor-Gene Q (QIAGEN); cycling conditions were as follows: 2 min at 50 °C, 10 min at 95 °C, followed by 40 cycles of 15 s at 95 °C, 30 s at 60 °C, and 30 s at 72 °C. Relative expression of *tcf21* was determined using the 2^−ΔΔCt^ method (Livak & Schmittgen, 2001) using *eef1a1l1 (ef1a)* as a housekeeping gene control. Sequences of primers are listed in Table S1.

### qPCR

gDNA was extracted from the lower phenol-chloroform phase and interphase remaining after RNA extraction using TRI Reagent (Sigma-Aldrich, T9424) according to the manufacturers’ recommendations. qPCR on gDNA was performed using 10 ng of template, 0.15 µM of forward and reverse primers, and Maxima SYBR Green qPCR Master Mix (2x) without ROX passive dye (Thermo Scientific, K0253) on Rotor-Gene Q (QIAGEN) under the same conditions as described above. Fold change of the *tcf21* gene was determined using the 2^−ΔΔCt^ method (Livak & Schmittgen, 2001). The *thyroglobulin precursor* (*TG*) gene was used as a reference locus (Kalvaityte et al., 2024; D. Wang et al., 2007). Sequences of primers are listed in Table S1.

### AFOG staining

Adult zebrafish hearts were harvested, fixed with 4% PFA for 2 h at RT on a nutator, cryopreserved in 30% sucrose, embedded into OCT Embedding Matrix (CellPath), and stored at -80 °C as described before (González-Rosa & Mercader, 2012). Hearts were sectioned into 10 µm sections using Cryotome (Thermo Scientific) and stored at -20 °C. AFOG (Acid Fuchsin-Orange G) staining was performed using an A.F.O.G. kit (BioGnost, AFOG-K-100) according to the manufacturer’s instructions. Imaging of AFOG-stained sections was performed using a Leica DM5500 B microscope with an HC PLAN APO 20x/0.70 objective. 60 dpci heart sections exhibiting the largest injury area from each heart were evaluated by an independent expert and scored blindly as completely regenerated, partially regenerated, and not regenerated.

### Immunostaining

Adult zebrafish hearts were harvested, fixed with 4% PFA for 2 h at RT on a nutator, cryopreserved in 30% sucrose, embedded into OCT Embedding Matrix (CellPath), and stored at -80 °C before sectioning (González-Rosa & Mercader, 2012). 8-10 µm sections were used for immunostaining as described before (Beisaw et al., 2020). Briefly, slides were incubated in sodium citrate buffer (10 mM Tri-sodium citrate, 0.05% Tween-20, pH 6) at 95°C for 45 min, washed twice in 0.1% Triton X-100 in PBS (PBSTr), twice in dH_2_O, and permeabilized in 3% H_2_O_2_ in methanol for 1 h at RT. Then, sections were washed twice in dH20, twice in PBSTr, and incubated in blocking solution (1x PBS, 2% FBS, 0.2% Triton X-100, 1% DMSO) for 2 h at RT. Primary antibodies were incubated overnight at 4°C, followed by three washes with PBSTr. Slides were incubated with secondary antibodies for 2 h at RT, washed three times with PBSTr, and mounted in a Mowiol mounting medium. Primary antibodies used in this study: anti-Aldh1a2 (GeneTex, GTX124302) at 1:500, anti-α-Actinin/ACTN1 (abcam, ab210557) at 1:200, embCMHC (DSHB, N2.261) at 5 μg/ml, anti-PCNA (Sigma-Aldrich, P8825) at 1:3000, and anti-MEF2A+MEF2C (abcam, ab197070) at 1:200. Alexa Fluor 488 (Invitrogen, A11001) and Alexa Fluor 594 (Invitrogen, A11012) secondary antibodies were used at 1:500. Imaging of immunostained sections was performed using a Leica TCS SP8 confocal scanning microscope with an HC PL APO 20x/0,75 CS2 objective. All measurements were done in at least three nonconsecutive sections exhibiting the largest injury area from each heart using ImageJ (v1.54f) (Schindelin et al., 2012).

### RNA-seq and data analysis

Adult *tcf21^flox/flox^, ubb^R^:CreER^T2^*, tcf21:DsRed2* fish were treated with 4-HT as described above. 2-3 ventricles per sample were collected at 0 dpt, 3 dpci, or 3 dps and used to extract RNA using TRI reagent (Sigma-Aldrich, T9424) according to the manufacturer’s recommendations. Untreated adult *tcf21^flox/flox^, ubb^R^:CreER^T2^*, tcf21:DsRed2* fish ventricles, collected at day 0, 3 dpci, or 3 dps, were used as a control. Extracted RNA was treated with DNase I (Thermo Scientific, 89836). mRNA purification, library preparation, and sequencing were carried out following standard Illumina protocols by Novogene Co., Ltd. (Cambridge, UK). Briefly, mRNA was purified using poly-T oligo-attached magnetic beads and fragmented. The first strand cDNA was synthesized using random hexamer primers, followed by the second strand cDNA synthesis. The library was quantified with Qubit and real-time PCR, and size distribution was detected with a bioanalyzer. Quantified libraries were pooled and sequenced on an Illumina NovaSeq 6000 S4 platform.

Raw reads were quality trimmed using TrimGalore (v. 0.6.6) (Krueger et al., 2023) and mapped to the zebrafish reference genome (GRCz11) using HISAT2 aligner (v2.1.0) (Kim et al., 2019). Reads were summarized on protein-coding gene level using StringTie (v2.1.1) (Pertea et al., 2016). Differential gene expression analysis was performed using the DESeq2 package (v1.38.3) (Love et al., 2014). Principal component analysis (PCA) was performed on VST-transformed data in R (v4.2.2) using DESeq2 (v1.38.3). Normalized counts were used to calculate the Z-score and draw a heatmap using ComplexHeatmap (v2.14.0). Gene ontology (GO) enrichment analysis of differentially expressed genes with an adjusted p-value below 0.05 was done using the DAVID database (v2024q4) (Huang et al., 2009; Sherman et al., 2022).

### Statistical analysis

All statistical analysis was performed with R software (v4.2.2). The Shapiro-Wilk test was used to check for the normality of the data. Comparative statistics between two sample groups were performed using the unpaired t-test for parametric data or the Wilcoxon rank-sum test for nonparametric data. The distribution of regeneration phenotype at 60 dpci was evaluated by Fisher’s Exact Test. Comparative statistics between more than two sample groups were performed using one-way ANOVA followed by Tukey’s post-hoc test for parametric data or the pairwise Wilcoxon rank-sum test with Bonferroni correction for nonparametric data.

## Data availability

RNA-seq data from this study have been deposited in the Gene Expression Omnibus (GEO) database under the accession number GSE289891.

## Acknowledgments

We sincerely thank João Cardeira-da-Silva (Inserm, France) for the advice on immunostaining heart sections.

---

**Fig. S1.**
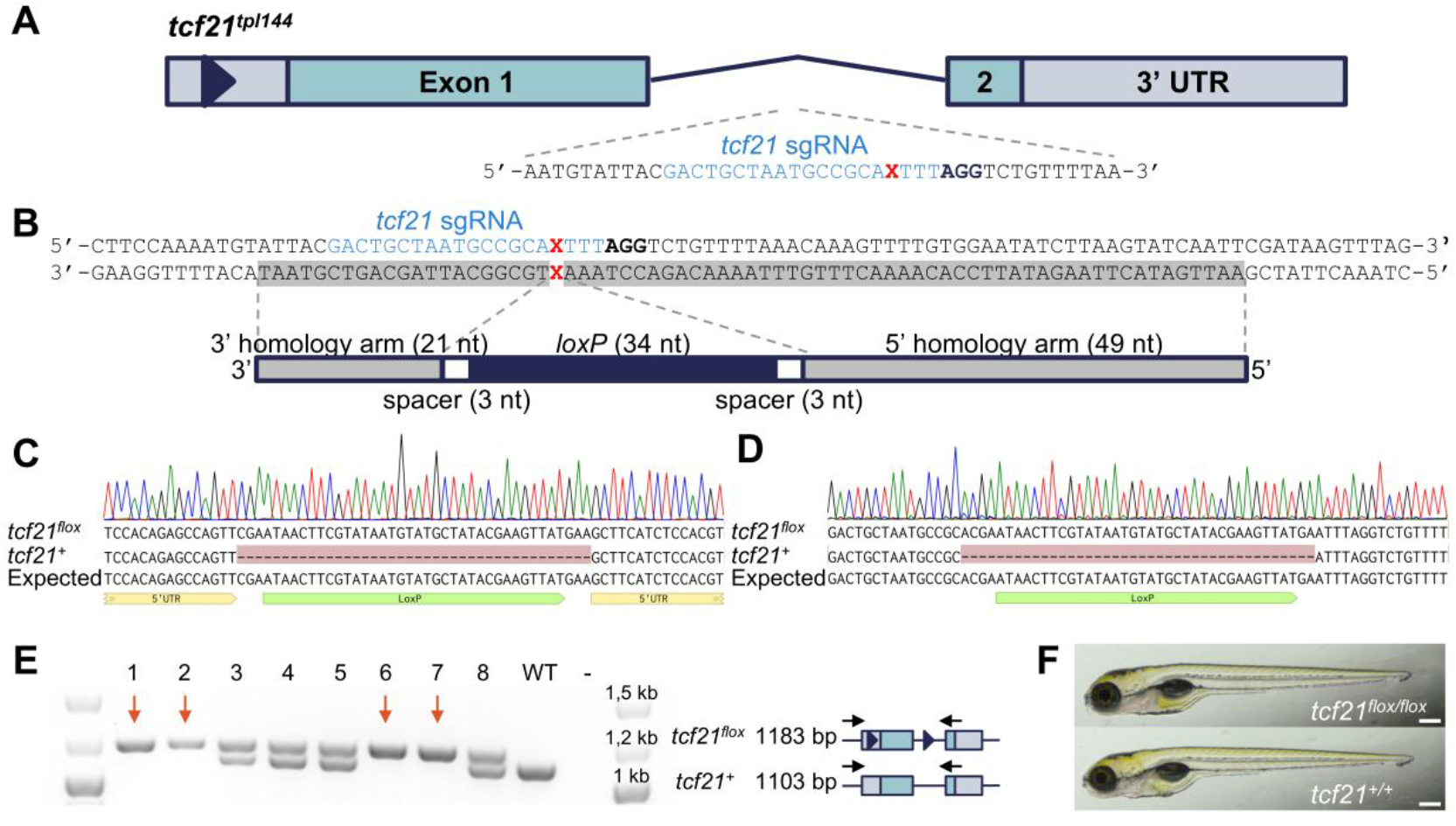
Integration of *loxP* site into the intron of *tcf21* gene. **A.** Diagram of the *tcf21^tpl144^* allele. Triangle marks *loxP* site. Sequence corresponding to sgRNA used to induce double-stranded break at first intron of *tcf21* gene is shown in blue, PAM motif in bold, and expected Cas9 cut site is indicated by a red X. **B.** Integration of *loxP* site into the intron of *tcf21*. ssODN with asymmetric homology arms was used to induce homology-directed repair. ssODN was in antisense strand to the PAM site (in bold) and composed of *loxP* site (dark blue) flanked by 3 nucleotide (nt) spacer sequence (white), 21 nt long 3’ homology arm, and 49 nt long 5’ homology arm. **C-D.** Sequence of the recovered *tcf21^flox^* allele containing perfect integration of the *loxP* sites at 5’ UTR **(C)** and first intron **(D)** of the *tcf21* gene. **E.** Offspring of heterozygous F1 fish incross were tail clipped and genotyped. Red arrows indicate fish homozygous for the *tcf21^flox^* allele. “WT” – *tcf21^+/+^* control; “-” – no template control (NTC). **F.** Bright field images of homozygous *tcf21^flox^* and wild-type 5 dpf larvae. Scale bar: 250 μm.

**Fig. S2.**
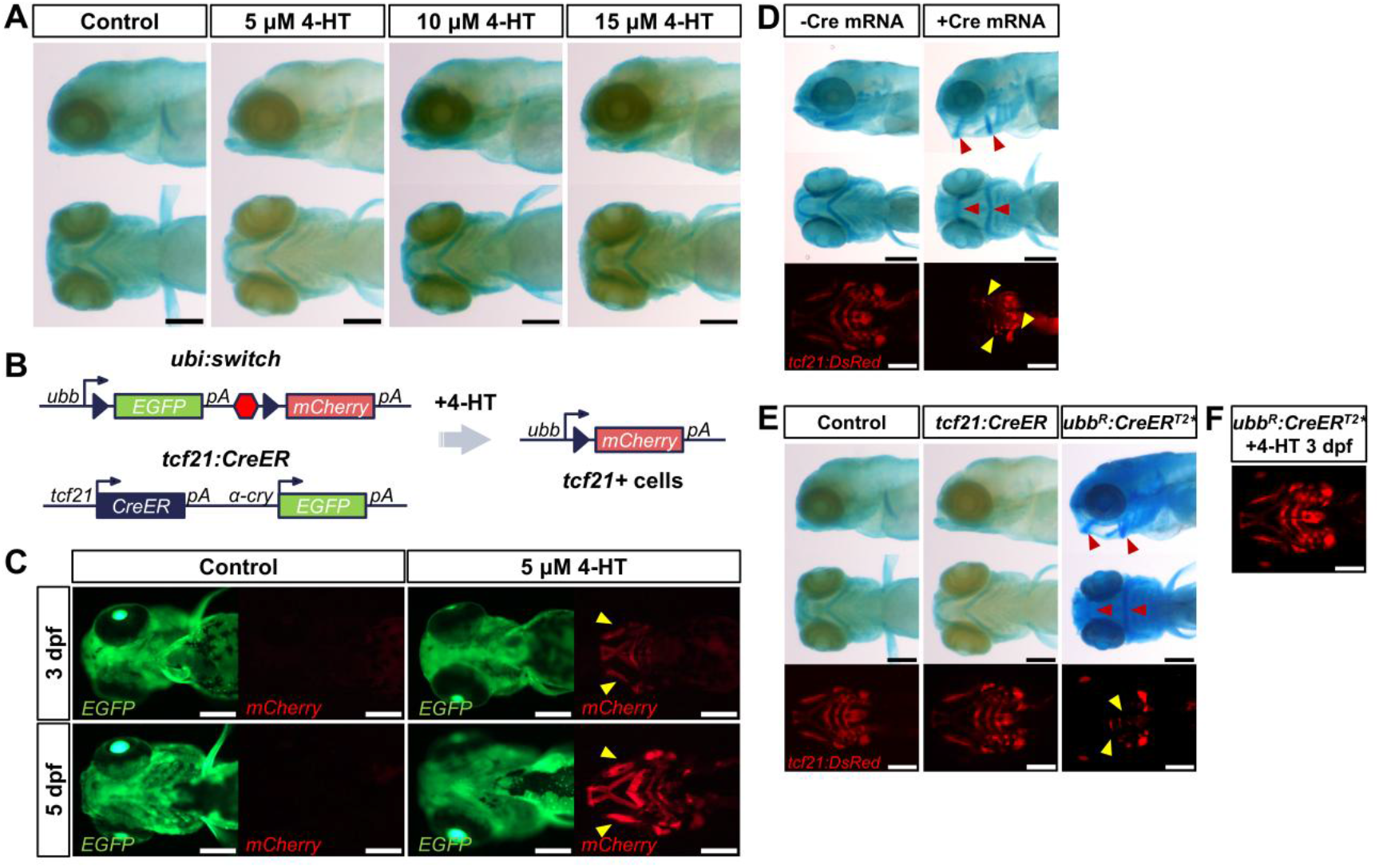
*tcf21:CreER* transgene does not induce loss-of-function phenotype in *tcf21^flox^* larvae. **A.** Alcian blue-stained 5dpf *tcf21^flox/flox^, tcf21:CreER* larvae treated with different 4-HT concentrations at 6 hpf. Control – *tcf21^flox/flox^* untreated larvae. **B.** Scheme illustrating *ubi:switch* and *tcf21:CreER* transgenes and the recombination outcome after 4-HT treatment. “pA” – polyA; red hexagon – transcription termination signal; triangles – *loxP* sites. **C.** *tcf21:CreER, ubi:switch* embryos were treated with 5 μM 4-HT from 6 hpf to 3 dpf and imaged at 3 and 5 dpf. Yellow arrowheads indicate successful recombination of *ubi:switch* transgene in *tcf21* expressing cells. Control embryos were not treated with 4-HT. **D.** Bright field images of alcian-blue-stained and fluorescence images of *tcf21^flox/flox^, tcf21:DsRed2* 5 dpf larvae injected or not-injected with Cre recombinase mRNA. **E.** Bright field of alcian-blue-stained and fluorescence images of *tcf21^flox/flox^, tcf21:DsRed2, tcf21:CreER* and *tcf21^flox/flox^, tcf21:DsRed2, ubb^R^:CreER^T2^\** 5 dpf larvae treated with 5 μM 4-HT from 6 hpf to 3 dpf. Control -*tcf21^flox/flox^, tcf21:DsRed2* larvae treated with 5 μM 4-HT from 6 hpf to 3 dpf. Red arrowheads in (**D, E**) mark defects in branchial arch formation; yellow arrowheads indicate loss of *tcf21:DsRed2* expression. **F.** Fluorescence image of 5dpf *tcf21^flox/flox^, tcf21:DsRed2, ubb^R^:CreER^T2^\** larvae treated with 5 μM 4-HT from 3 dpf to 5 dpf. Scale bar: 200 μm in (**A,C-F**).

**Fig. S3.**
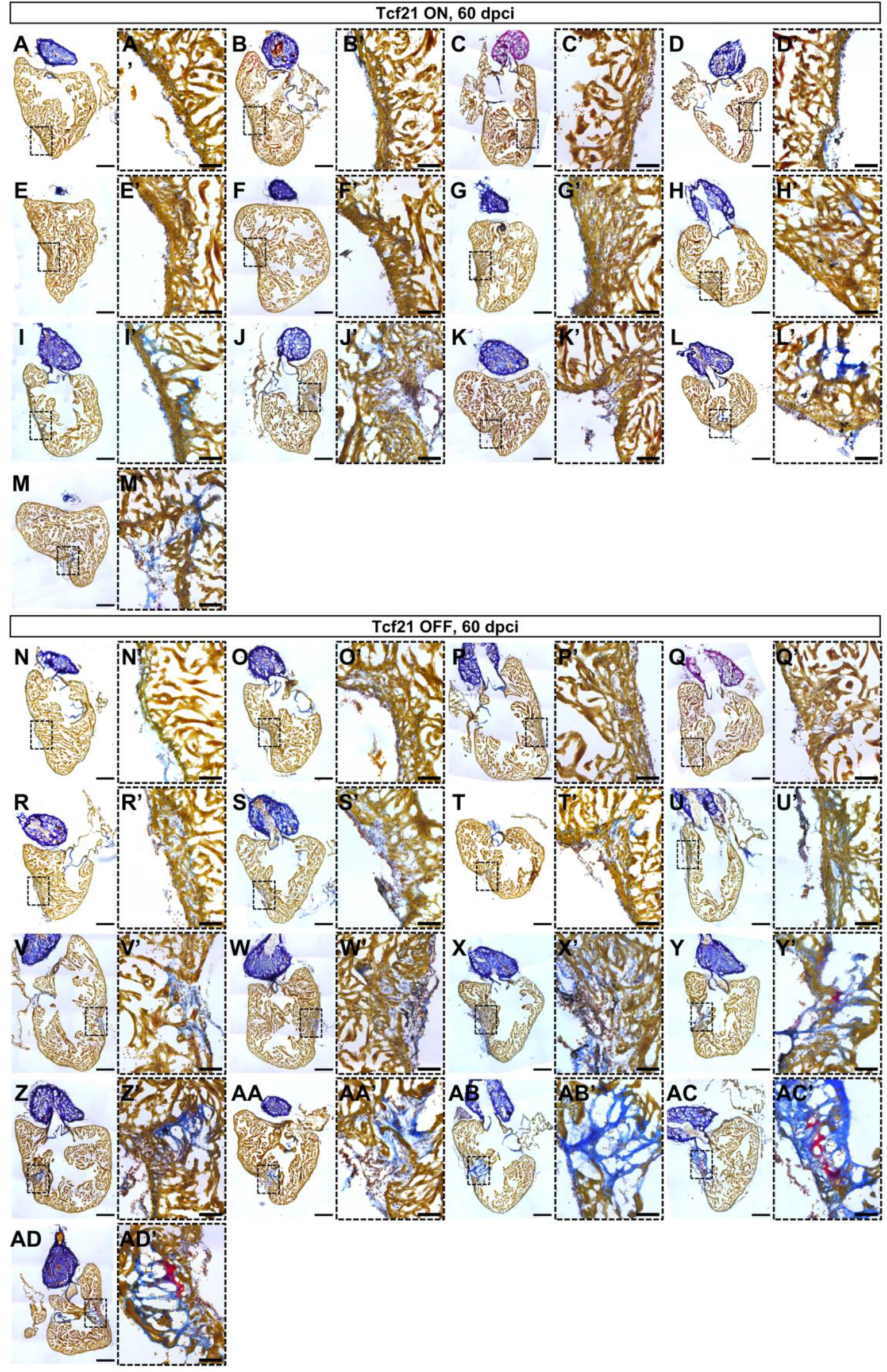
Conditional *tcf21^flox^* knock-out in adult fish impairs heart regeneration. 60 dpci heart sections exhibiting the largest injury area of (**A-M**) Tcf21 ON, and (**N-AD**) Tcf21 OFF fish. Muscle tissue is visualized in brown, fibrin in red and collagen in blue. Representative sections of the hearts (scale bar: 200 μm) with higher magnification images of the marked area (scale bar: 50 μm) are presented.

**Fig. S4.**
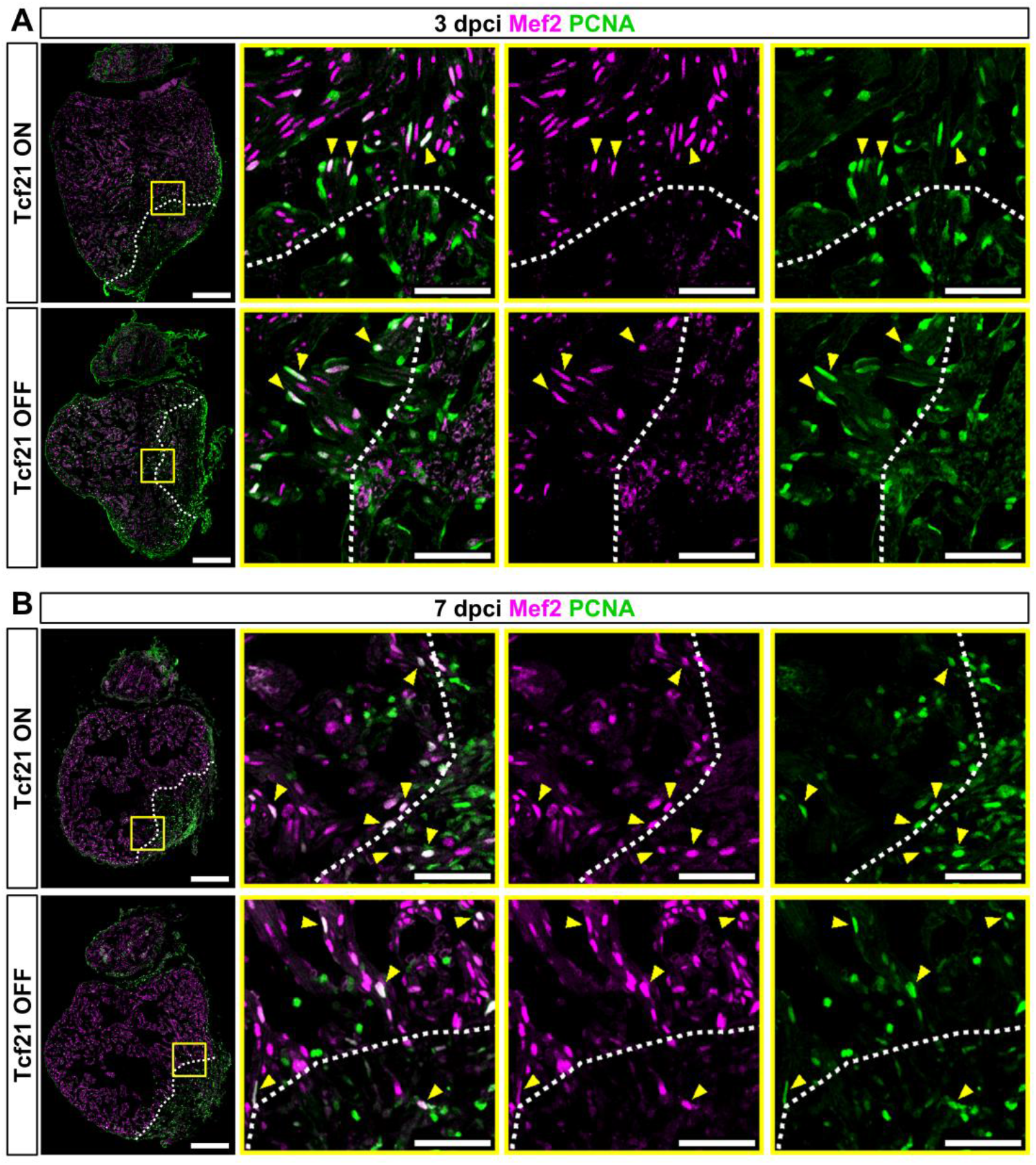
Cardiomyocyte (CM) proliferation is not affected by the loss of Tcf21. **A-B.** PCNA and Mef2 staining of Tcf21 ON and Tcf21 OFF fish heart sections at 3 dpci (**A**) and 7 dpci (**B**) (scale bar: 200 μm). White dashed lines indicate wound border, yellow arrowheads point to proliferating (PCNA+) CMs, and yellow boxes mark the areas shown in zoomed images (scale bar: 50 μm).

**Table S1.**
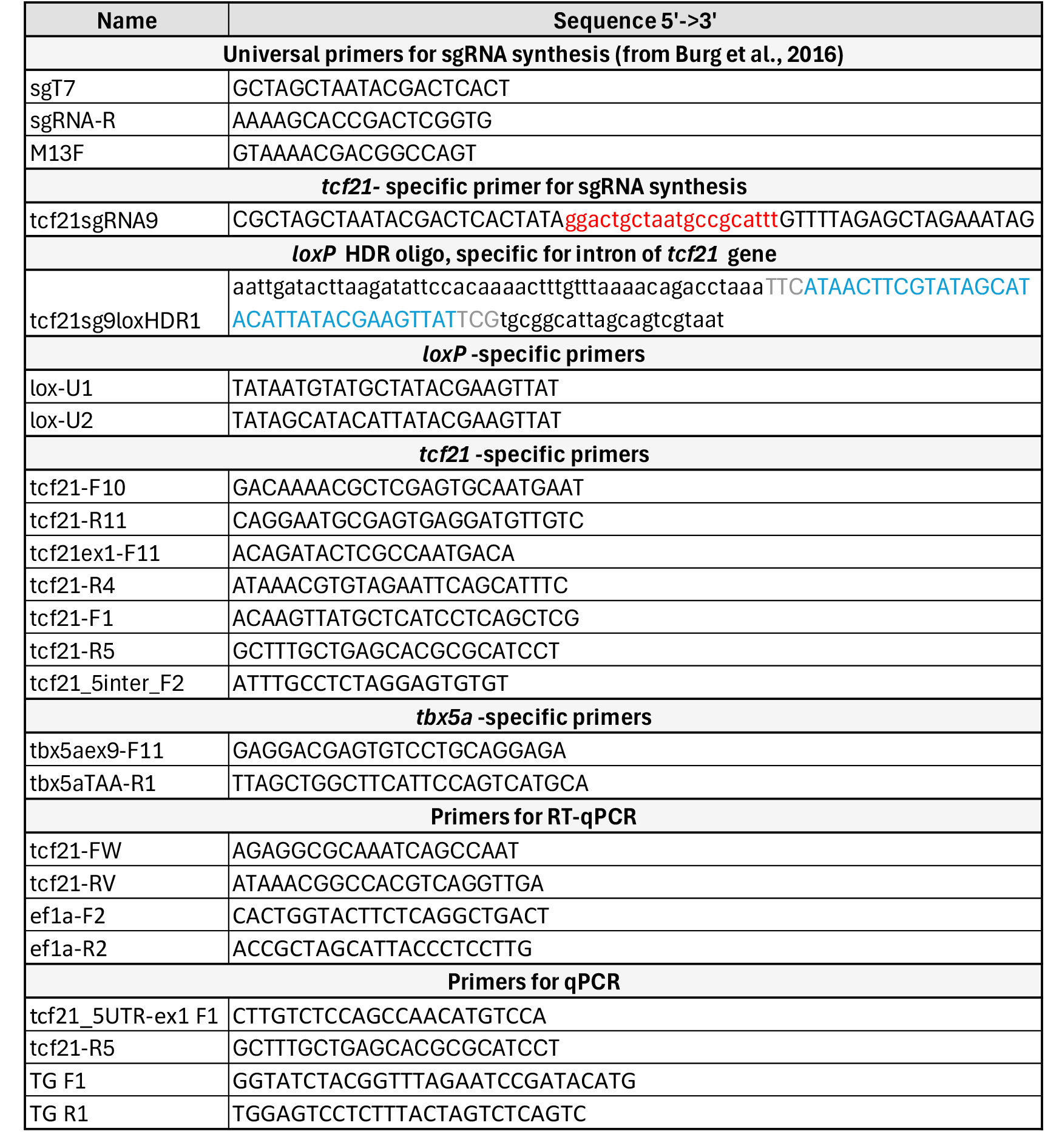
Primers and oligonucleotides used in this study. The *tcf21* target sequence of sgRNA is marked in red, the *loxP* site in ssODN is in blue, and spacer sequences are in grey.

## REFERENCES

Acharya, A., Baek, S. T., Huang, G., Eskiocak, B., Goetsch, S., Sung, C. Y., Banfi, S., Sauer, M. F., Olsen, G. S., Duffield, J. S., Olson, E. N., & Tallquist, M. D. (2012). The bHLH transcription factor Tcf21 is required for lineage-specific EMT of cardiac fibroblast progenitors. Development (Cambridge, England), 139(12), 2139–2149. 10.1242/dev.079970

Allanki, S., Strilic, B., Scheinberger, L., Onderwater, Y. L., Marks, A., Günther, S., Preussner, J., Kikhi, K., Looso, M., Stainier, D. Y. R., & Reischauer, S. (2021). Interleukin-11 signaling promotes cellular reprogramming and limits fibrotic scarring during tissue regeneration. Science Advances, 7(37). 10.1126/sciadv.abg6497

Angom, R. S., Wang, Y., Wang, E., Dutta, S. K., & Mukhopadhyay, D. (2023). tConditional, Tissue-Specific CRISPR/Cas9 Vector System in Zebrafish Reveals the Role of Nrp1b in Heart Regeneration. Arteriosclerosis, Thrombosis, and Vascular Biology, 43(10), 1921–1934. 10.1161/ATVBAHA.123.319189

Bakūnaitė, E., Gečaitė, E., Lazutka, J., & Balciunas, D. (2024). Highly efficient tamoxifen-inducible Cre recombination in embryonic, larval and adult zebrafish. BioRxiv. 10.1101/2024.03.21.586128

Balciuniene, J., & Balciunas, D. (2013). Gene Trapping Using Gal4 in Zebrafish. JoVE, 79, e50113. doi:10.3791/50113

Balciuniene, J., Nagelberg, D., Walsh, K. T., Camerota, D., Georlette, D., Biemar, F., Bellipanni, G., & Balciunas, D. (2013). Efficient disruption of Zebrafish genes using a Gal4-containing gene trap. BMC Genomics, 14(1), 619. 10.1186/1471-2164-14-619

Beisaw, A., Kuenne, C., Guenther, S., Dallmann, J., Wu, C. C., Bentsen, M., Looso, M., & Stainier, D. Y. R. (2020). AP-1 Contributes to Chromatin Accessibility to Promote Sarcomere Disassembly and Cardiomyocyte Protrusion during Zebrafish Heart Regeneration. Circulation Research, 126(12), 1760–1778. 10.1161/CIRCRESAHA.119.316167

Ben-Yair, R., Butty, V. L., Busby, M., Qiu, Y., Levine, S. S., Goren, A., Boyer, L. A., Geoffrey Burns, C., & Burns, C. E. (2019). H3K27me3-mediated silencing of structural genes is required for zebrafish heart regeneration. Development (Cambridge), 146(19). 10.1242/dev.178632

Bergmann, O., Bhardwaj, R. D., Bernard, S., Zdunek, S., Barnabé-Heide, F., Walsh, S., Zupicich, J., Alkass, K., Buchholz, B. A., Druid, H., Jovinge, S., & Frisén, J. (2009). Evidence for cardiomyocyte renewal in humans. Science, 324(5923), 98–102. 10.1126/science.1164680

Boezio, G. L. M., Zhao, S., Gollin, J., Priya, R., Mansingh, S., Guenther, S., Fukuda, N., Gunawan, F., & Stainier, D. Y. R. (2023). The developing epicardium regulates cardiac chamber morphogenesis by promoting cardiomyocyte growth. DMM Disease Models and Mechanisms, 16(5). 10.1242/dmm.049571

Braitsch, C. M., Combs, M. D., Quaggin, S. E., & Yutzey, K. E. (2012). Pod1/Tcf21 is regulated by retinoic acid signaling and inhibits differentiation of epicardium-derived cells into smooth muscle in the developing heart. Developmental Biology, 368(2), 345–357. 10.1016/j.ydbio.2012.06.002

Braitsch, C. M., Kanisicak, O., van Berlo, J. H., Molkentin, J. D., & Yutzey, K. E. (2013). Differential expression of embryonic epicardial progenitor markers and localization of cardiac fibrosis in adult ischemic injury and hypertensive heart disease. Journal of Molecular and Cellular Cardiology, 65, 108–119. 10.1016/j.yjmcc.2013.10.005

Broughton, K. M., Wang, B. J., Firouzi, F., Khalafalla, F., Dimmeler, S., Fernandez-Aviles, F., & Sussman, M. A. (2018). Mechanisms of Cardiac Repair and Regeneration. Circ Res., 122(8), 1151–1163. doi:10.1161/CIRCRESAHA.117.312586

Burg, L., Palmer, N., Kikhi, K., Miroshnik, E. S., Rueckert, H., Gaddy, E., Cunningham, C. M., Mattonet, K., Lai, S., Marin-Juez, R., Waring, R. B., Stainier, D. Y. R., & Balciunas, D. (2018). Conditional mutagenesis by oligonucleotide-mediated integration of loxP sites in zebrafish. PLOS Genetics, 1–26. 10.1371/journal.pgen.1007754

Burg, L., Zhang, K., Bonawitz, T., Grajevskaja, V., Bellipanni, G., Waring, R., & Balciunas, D. (2016). Internal epitope tagging informed by relative lack of sequence conservation. Scientific Reports, 6, 1–8. 10.1038/srep36986

Cao, J., & Poss, K. D. (2018). The epicardium as a hub for heart regeneration. Nat Rev Cardiol., 15(10), 631–647. 10.1038/s41569-018-0046-4

Chablais, F., Veit, J., Rainer, G., & Jawiska, A. (2011). The zebrafish heart regenerates after cryoinjury-induced myocardial infarction. BMC Developmental Biology, 11. 10.1186/1471-213X-11-21

Constanty, F., Wu, B., Wei, K. H., Lin, I. T., Dallmann, J., Guenther, S., Lautenschlaeger, T., Priya, R., Lai, S. L., Stainier, D. Y. R., & Beisaw, A. (2024). Border-zone cardiomyocytes and macrophages contribute to remodeling of the extracellular matrix to promote cardiomyocyte invasion during zebrafish cardiac regeneration. BioRxiv.

Fang, Y., Gupta, V., Karra, R., Holdway, J. E., Kikuchi, K., & Poss, K. D. (2013). Translational profiling of cardiomyocytes identifies an early Jak1/Stat3 injury response required for zebrafish heart regeneration. Proceedings of the National Academy of Sciences of the United States of America, 110(33), 13416–13421. 10.1073/pnas.1309810110

Felker, A., Nieuwenhuize, S., Dolbois, A., Blazkova, K., Hess, C., Low, L. W. L., Burger, S., Samson, N., Carney, T. J., Bartunek, P., Nevado, C., & Mosimann, C. (2016). In vivo performance and properties of Tamoxifen metabolites for CreERT2 control. PLoS ONE, 11(4), 1–17. 10.1371/journal.pone.0152989

Gebauer, J. M., Kobbe, B., Paulsson, M., & Wagener, R. (2016). tStructure, evolution and expression of collagen XXVIII: Lessons from the zebrafish. Matrix Biology, 49, 106–119. 10.1016/j.matbio.2015.07.001

Gemberling, M., Karra, R., Dickson, A. L., & Poss, K. D. (2015). Nrg1 is an injury-induced cardiomyocyte mitogen for the endogenous heart regeneration program in zebrafish. ELife, 2015(4), 1–17. 10.7554/eLife.05871

González-Rosa, J. M., Martín, V., Peralta, M., Torres, M., & Mercader, N. (2011). Extensive scar formation and regression during heart regeneration after cryoinjury in zebrafish. Development, 138(9), 1663–1674. 10.1242/dev.060897

González-Rosa, J. M., & Mercader, N. (2012). Cryoinjury as a myocardial infarction model for the study of cardiac regeneration in the zebrafish. Nature Protocols, 7(4), 782–788. 10.1038/nprot.2012.025

Grajevskaja, V., Camerota, D., Bellipanni, G., Balciuniene, J., & Balciunas, D. (2018). Analysis of a conditional gene trap reveals that tbx5a is required for heart regeneration in zebrafish. PLoS ONE, 13(6), 1–14. 10.1371/journal.pone.0197293

Housley, M. P., Njaine, B., Ricciardi, F., Stone, O. A., Hölper, S., Krüger, M., Kostin, S., & Stainier, D. Y. R. (2016). Cavin4b/Murcb Is Required for Skeletal Muscle Development and Function in Zebrafish. PLoS Genetics, 12(6), 1–19. 10.1371/journal.pgen.1006099

Itou, J., Oishi, I., Kawakami, H., Glass, T. J., Richter, J., Johnson, A., Lund, T. C., & Kawakami, Y. (2012). Migration of cardiomyocytes is essential for heart regeneration in zebrafish. Development (Cambridge), 139(22), 4133–4142. 10.1242/dev.079756

Jao, L. E., Wente, S. R., & Chen, W. (2013). Efficient multiplex biallelic zebrafish genome editing using a CRISPR nuclease system. PNAS, 110(34), 13904–13909. 10.1073/pnas.1308335110

Jayaraj, J. C., Davatyan, K., Subramanian, S. S., & Priya, J. (2018). Epidemiology of Myocardial Infarction. In B. Pamukçu (Ed.), Myocardial Infarction. IntechOpen. 10.5772/intechopen.74768

Johansen, A. K. Z., Kasam, R. K., Vagnozzi, R. J., Lin, S. C. J., Gomez-Arroyo, J. G., Shittu, A., Bowers, S. L. K., Kuwabara, Y., Grimes, K. M., Warrick, K., Sargent, M. A., Baldwin, T. A., Quaggin, S. E., Barski, A., & Molkentin, J. D. (2025). Transcription Factor 21 Regulates Cardiac Myofibroblast Formation and Fibrosis. Circulation Research, 136, 44–58. 10.1161/CIRCRESAHA.124.325527

Jopling, C., Sleep, E., Raya, M., Martí, M., Raya, A., & Belmonte, J. C. I. (2010). Zebrafish heart regeneration occurs by cardiomyocyte dedifferentiation and proliferation. Nature, 464(7288), 606–609. 10.1038/nature08899

Kalvaitytė, M., & Balciunas, D. (2022). Conditional mutagenesis strategies in zebrafish. Trends in Genetics, 1–13. 10.1016/j.tig.2022.04.007

Kalvaityte, M., Gabrilaviciute, S., & Balciunas, D. (2024). Rapid generation of single-insertion transgenics by Tol2 transposition in zebrafish. Developmental Dynamics, May, 1–10. 10.1002/dvdy.719

Kanisicak, O., Khalil, H., Ivey, M. J., Karch, J., Maliken, B. D., Correll, R. N., Brody, M. J., Lin, S. C. J., Aronow, B. J., Tallquist, M. D., & Molkentin, J. D. (2016). Genetic lineage tracing defines myofibroblast origin and function in the injured heart. Nature Communications, 7. 10.1038/ncomms12260

Kikuchi, K., Gupta, V., Wang, J., Holdway, J. E., Wills, A. A., Fang, Y., & Poss, K. D. (2011). Tcf21+ epicardial cells adopt non-myocardial fates during zebrafish heart development and regeneration. Development, 138(14), 2895–2902. 10.1242/dev.067041

Kikuchi, K., Holdway, J. E., Major, R. J., Blum, N., Dahn, R. D., Begemann, G., & Poss, K. D. (2011). Retinoic Acid Production by Endocardium and Epicardium Is an Injury Response Essential for Zebrafish Heart Regeneration. Developmental Cell, 20(3), 397–404. 10.1016/j.devcel.2011.01.010

Kikuchi, K., Holdway, J. E., Werdich, A. A., Anderson, R. M., Fang, Y., Egnaczyk, G. F., Evans, T., MacRae, C. A., Stainier, D. Y. R., & Poss, K. D. (2010). Primary contribution to zebrafish heart regeneration by gata4+ cardiomyocytes. Nature, 464(7288), 601–605. 10.1038/nature08804

Kim, D., Paggi, J. M., Park, C., Bennett, C., & Salzberg, S. L. (2019). Graph-based genome alignment and genotyping with HISAT2 and HISAT-genotype. Nature Biotechnology, 37(8), 907–915. 10.1038/s41587-019-0201-4

Krueger, F., James, F., Ewels, P., Afyounian, E., Weinstein, M., Schuster-Boeckler, B., Hulselmans, G., & sclamons. (2023). FelixKrueger/TrimGalore: v0.6.10 - add default decompression path. Zenodo. 10.5281/zenodo.7598955

Lee, G. H., Chang, M. Y., Hsu, C. H., & Chen, Y. H. (2011). Essential roles of basic helix-loop-helix transcription factors, Capsulin and Musculin, during craniofacial myogenesis of zebrafish. Cellular and Molecular Life Sciences, 68(24), 4065–4078. 10.1007/s00018-011-0637-2

Lepilina, A., Coon, A. N., Kikuchi, K., Holdway, J. E., Roberts, R. W., Burns, C. G., & Poss, K. D. (2006). A Dynamic Epicardial Injury Response Supports Progenitor Cell Activity during Zebrafish Heart Regeneration. Cell, 127, 607–619. 10.1016/j.cell.2006.08.052

Livak, K. J., & Schmittgen, T. D. (2001). Analysis of relative gene expression data using real-time quantitative PCR and the 2-ΔΔCT method. Methods, 25(4), 402–408. 10.1006/meth.2001.1262

Lodrini, A. M., & Goumans, M. J. (2021). Cardiomyocytes Cellular Phenotypes After Myocardial Infarction. Frontiers in Cardiovascular Medicine, 8(November), 1–11. 10.3389/fcvm.2021.750510

Lu, J., Chang, P., Richardson, J. A., Gan, L., Weiler, H., & Olson, E. N. (2000). The basic helix-loop-helix transcription factor capsulin controls spleen organogenesis. Proceedings of the National Academy of Sciences of the United States of America, 97(17), 9525–9530. 10.1073/pnas.97.17.9525

Lu, J., Richardson, J. A., & Olson, E. N. (1998). Capsulin : a novel bHLH transcription factor expressed in epicardial progenitors and mesenchyme of visceral organs. Mechanisms of Development, 73, 23–32.

Marín-Juez, R., El-Sammak, H., Helker, C. S. M., Kamezaki, A., Mullapuli, S. T., Bibli, S. I., Foglia, M. J., Fleming, I., Poss, K. D., & Stainier, D. Y. R. (2019). Coronary Revascularization During Heart Regeneration Is Regulated by Epicardial and Endocardial Cues and Forms a Scaffold for Cardiomyocyte Repopulation. Developmental Cell, 51(4), 503–515.e4. 10.1016/j.devcel.2019.10.019

Martin, S. S., Aday, A. W., Almarzooq, Z. I., Anderson, C. A. M., Arora, P., Avery, C. L., Baker-Smith, C. M., Gibbs, B. B., Beaton, A. Z., Boehme, A. K., Commodore-Mensah, Y., Currie, M. E., Elkind, M. S. V, Evenson, K. R., Generoso, G., Heard, D. G., Hiremath, S., Johansen, M. C., Kalani, R., … Subcommittee, S. S. (2024). 2024 Heart Disease and Stroke Statistics: A Report of US and Global Data From the American Heart Association. Circulation, 149(8), e347–e913. 10.1161/CIR.0000000000001209

Morikawa, Y., Zhang, M., Heallen, T., Leach, J., Tao, G., Xiao, Y., Bai, Y., Li, W., Willerson, J. T., & Martin, J. F. (2015). Actin cytoskeletal remodeling with protrusion formation is essential for heart regeneration in Hippo-deficient mice. Science Signaling, 8(375), 1–13. 10.1126/scisignal.2005781

Mosimann, C., Kaufman, C. K., Li, P., Pugach, E. K., Tamplin, O. J., & Zon, L. I. (2011). Ubiquitous transgene expression and Cre-based recombination driven by the ubiquitin promoter in zebrafish. Development, 138(1), 169–177. 10.1242/dev.059345

Münch, J., Grivas, D., González-Rajal, Á., Torregrosa-Carrión, R., & de la Pompa, J. L. (2017). Notch signalling restricts inflammation and Serpine1 expression in the dynamic endocardium of the regenerating zebrafish heart. Development (Cambridge), 144(8), 1425–1440. 10.1242/dev.143362

Nagelberg, D., Wang, J., Su, R., Torres-Vázquez, J., Targoff, K. L., Poss, K. D., & Knaut, H. (2015). tOrigin, specification, and plasticity of the great vessels of the heart. Current Biology, 25(16), 2099–2110. 10.1016/j.cub.2015.06.076

Ogawa, M., Geng, F.-S., Humphreys, D. T., Kristianto, E., Sheng, D. Z., Hui, S. P., Zhang, Y., Sugimoto, K., Nakayama, M., Zheng, D., Hesselson, D., Hodson, M. P., Bogdanovic, O., & Kikuchi, K. (2021). Krüppel-like factor 1 is a core cardiomyogenic trigger in zebrafish. Science, 372(6538), 201–205. 10.1126/science.abe2762

Pertea, M., Kim, D., Pertea, G. M., Leek, J. T., & Salzberg, S. L. (2016). Transcript-level expression analysis of RNA-seq experiments with HISAT, StringTie and Ballgown. Nature Protocols, 11(9), 1650–1667. 10.1038/nprot.2016.095

Pfefferli, C., & Jaźwińska, A. (2017). The careg element reveals a common regulation of regeneration in the zebrafish myocardium and fin. Nature Communications, 8(May). 10.1038/ncomms15151

Poss, K. D., Wilson, L. G., & Keating, M. T. (2002). Heart Regeneration in Zebrafish. Science, 298(5601), 2188–2191. 10.1126/science.1077857

Quaggin, S. E., Schwartz, L., Cui, S., Igarashi, P., Deimling, J., Post, M., & Rossant, J. (1999). The basic-helix-loop-helix protein Pod1 is critically important for kidney and lung organogenesis. Development, 126(24), 5771–5783. 10.1242/dev.126.24.5771

Quijada, P., Trembley, M. A., & Small, E. M. (2020). The Role of the Epicardium during Heart Development and Repair. Circulation Research, 126, 377–394. 10.1161/CIRCRESAHA.119.315857

Rajan, A. R. D., Huang, Y., Stundl, J., Chu, K., Irodi, A., Yang, Z., Applegate, B. E., & Bronner, M. E. (2024). Generation of a zebrafish neurofibromatosis model via inducible knockout of nf2. DMM Disease Models and Mechanisms, 17. 10.1242/dmm.050862

Sallin, P., de Preux Charles, A. S., Duruz, V., Pfefferli, C., & Jaźwińska, A. (2015). A dual epimorphic and compensatory mode of heart regeneration in zebrafish. Developmental Biology, 399(1), 27–40. 10.1016/j.ydbio.2014.12.002

Sánchez-Iranzo, H., Galardi-Castilla, M., Minguillón, C., Sanz-Morejón, A., González-Rosa, J. M., Felker, A., Ernst, A., Guzmán-Martínez, G., Mosimann, C., & Mercader, N. (2018). Tbx5a lineage tracing shows cardiomyocyte plasticity during zebrafish heart regeneration. Nature Communications, 9(1). 10.1038/s41467-017-02650-6

Schindelin, J., Arganda-Carreras, I., Frise, E., Kaynig, V., Longair, M., Pietzsch, T., Preibisch, S., Rueden, C., Saalfeld, S., Schmid, B., Tinevez, J.-Y., White, D. J., Hartenstein, V., Eliceiri, K., Tomancak, P., & Cardona, A. (2012). Fiji: an open-source platform for biological-image analysis. Nature Methods, 9(7), 676–682. 10.1038/nmeth.2019

Schnabel, K., Wu, C. C., Kurth, T., & Weidinger, G. (2011). Regeneration of cryoinjury induced necrotic heart lesions in zebrafish is associated with epicardial activation and cardiomyocyte proliferation. PLoS ONE, 6(4). 10.1371/journal.pone.0018503

Shen, J., Cao, B., Wang, Y., Ma, C., Zeng, Z., Liu, L., Li, X., Tao, D., Gong, J., & Xie, D. (2018). Hippo component YAP promotes focal adhesion and tumour aggressiveness via transcriptionally activating THBS1/FAK signalling in breast cancer. Journal of Experimental and Clinical Cancer Research, 37(1), 1–17. 10.1186/s13046-018-0850-z

Simões, F. C., & Riley, P. R. (2018). The ontogeny, activation and function of the epicardium during heart development and regeneration. Development (Cambridge), 145(7). 10.1242/dev.155994

Sugimoto, K., Hui, S. P., Sheng, D. Z., & Kikuchi, K. (2017). Dissection of zebrafish shha function using site-specific targeting with a Cre-dependent genetic switch. ELife, 6, 1–20. 10.7554/eLife.24635

Sun, Y., Kiani, M. F., Postlethwaite, A. E., & Weber, K. T. (2002). Infarct scar as living tissue. Basic Research in Cardiology, 97(5), 343–347. 10.1007/s00395-002-0365-8

Tandon, P., Miteva, Y. V, Kuchenbrod, L. M., Cristea, I. M., & Conlon, F. L. (2013). Tcf21 regulates the specification and maturation of proepicardial cells. Development, 2421, 2409–2421. 10.1242/dev.093385

van Wijk, B., Gunst, Q. D., Moorman, A. F. M., & van den Hoff, M. J. B. (2012). Cardiac Regeneration from Activated Epicardium. PLoS ONE, 7(9). 10.1371/journal.pone.0044692

Vite, A., Li, J., & Radice, G. L. (2015). New functions for alpha-catenins in health and disease: from cancer to heart regeneration. Cell and Tissue Research, 360(3), 773–783. 10.1007/s00441-015-2123-x

Wang, D., Jao, L. E., Zheng, N., Dolan, K., Ivey, J., Zonies, S., Wu, X., Wu, K., Yang, H., Meng, Q., Zhu, Z., Zhang, B., Lin, S., & Burgess, S. M. (2007). Efficient genome-wide mutagenesis of zebrafish genes by retroviral insertions. Proceedings of the National Academy of Sciences of the United States of America, 104(30), 12428–12433. 10.1073/pnas.0705502104

Wang, J., Cao, J., Dickson, A. L., & Poss, K. D. (2015). Epicardial regeneration is guided by cardiac outflow tract and Hedgehog signalling. Nature, 522(7555), 226–230. 10.1038/nature14325

Wang, J., Karra, R., Dickson, A. L., & Poss, K. D. (2013). Fibronectin is deposited by injury-activated epicardial cells and is necessary for zebrafish heart regeneration. Developmental Biology, 382(2), 427–435. 10.1016/j.ydbio.2013.08.012

Wang, J., Panáková, D., Kikuchi, K., Holdway, J. E., Gemberling, M., Burris, J. S., Singh, S. P., Dickson, A. L., Lin, Y. F., Khaled Sabeh, M., Werdich, A. A., Yelon, D., MacRae, C. A., & Poss, K. D. (2011). The regenerative capacity of zebrafish reverses cardiac failure caused by genetic cardiomyocyte depletion. Development, 138(16), 3421–3430. 10.1242/dev.068601

Weinberger, M., Simões, F. C., Patient, R., Sauka-Spengler, T., & Riley, P. R. (2020). Functional Heterogeneity within the Developing Zebrafish Epicardium. Developmental Cell, 52(5), 574–590.e6. 10.1016/j.devcel.2020.01.023

Weinberger, M., Simões, F. C., Sauka-Spengler, T., & Riley, P. R. (2024). Distinct epicardial gene regulatory programmes drive development and regeneration of the zebrafish heart. Developmental Cell, 59, 1–17. 10.1016/j.devcel.2023.12.012

Wu, C.-C., Kruse, F., Vasudevarao, M. D., Junker, J. P., Zebrowski, D. C., Fischer, K., Noël, E. S., Grün, D., Berezikov, E., Engel, F. B., Van Oudenaarden, A., Weidinger, G., & Bakkers, J. (2016). Spatially Resolved Genome-wide Transcriptional Profiling Identifies BMP Signaling as Essential Regulator of Zebrafish Cardiomyocyte Regeneration. Developmental Cell, 36(1), 36–49. 10.1016/j.devcel.2015.12.010

Xia, Y., Duca, S., Perder, B., Dündar, F., Zumbo, P., Qiu, M., Yao, J., Cao, Y., Harrison, M. R. M., Zangi, L., Betel, D., & Cao, J. (2022). Activation of a transient progenitor state in the epicardium is required for zebrafish heart regeneration. Nature Communications, 13(1). 10.1038/s41467-022-35433-9

Zhang, R., Yang, J., Zhu, J., & Xu, X. (2009). Depletion of zebrafish Tcap leads to muscular dystrophy via disrupting sarcomere-membrane interaction, not sarcomere assembly. Human Molecular Genetics, 18(21), 4130–4140. 10.1093/hmg/ddp362

Zhou, Z., Zheng, L., Tang, C., Chen, Z., Zhu, R., Peng, X., Wu, X., & Zhu, P. (2020). Identification of Potentially Relevant Genes for Excessive Exercise-Induced Pathological Cardiac Hypertrophy in Zebrafish. Frontiers in Physiology, 11(November), 1–13. 10.3389/fphys.2020.565307

